# Deep convolutional models improve predictions of macaque V1 responses to natural images

**DOI:** 10.1101/201764

**Authors:** Santiago A. Cadena, George H. Denfield, Edgar Y. Walker, Leon A. Gatys, Andreas S. Tolias, Matthias Bethge, Alexander S. Ecker

## Abstract

Despite great efforts over several decades, our best models of primary visual cortex (V1) still predict spiking activity quite poorly when probed with natural stimuli, highlighting our limited understanding of the nonlinear computations in V1. Recently, two approaches based on deep learning have been successfully applied to neural data: On the one hand, transfer learning from networks trained on object recognition worked remarkably well for predicting neural responses in *higher areas* of the primate ventral stream, but has not yet been used to model spiking activity in early stages such as V1. On the other hand, data-driven models have been used to predict neural responses in the *early* visual system (retina and V1) of mice, but not primates. Here, we test the ability of both approaches to predict spiking activity in response to natural images in V1 of awake monkeys. Even though V1 is rather at an early to intermediate stage of the visual system, we found that the transfer learning approach performed similarly well to the data-driven approach and both outperformed classical linear-nonlinear and wavelet-based feature representations that build on existing theories of V1. Notably, transfer learning using a pre-trained feature space required substantially less experimental time to achieve the same performance. In conclusion, multi-layer convolutional neural networks (CNNs) set the new state of the art for predicting neural responses to natural images in primate V1 and deep features learned for object recognition are better explanations for V1 computation than all previous filter bank theories. This finding strengthens the necessity of V1 models that are multiple nonlinearities away from the image domain and it supports the idea of explaining early visual cortex based on high-level functional goals.

**Author summary:** Predicting the responses of sensory neurons to arbitrary natural stimuli is of major importance for understanding their function. Arguably the most studied cortical area is primary visual cortex (V1), where many models have been developed to explain its function. However, the most successful models built on neurophysiologists’ intuitions still fail to account for spiking responses to natural images. Here, we model spiking activity in primary visual cortex (V1) of monkeys using deep convolutional neural networks (CNNs), which have been successful in computer vision. We both trained CNNs directly to fit the data, and used CNNs trained to solve a high-level task (object categorization). With these approaches, we are able to outperform previous models and improve the state of the art in predicting the responses of early visual neurons to natural images. Our results have two important implications. First, since V1 is the result of several nonlinear stages, it should be modeled as such. Second, functional models of entire visual pathways, of which V1 is an early stage, do not only account for higher areas of such pathways, but also provide useful representations for V1 predictions.

## Introduction

An essential step towards understanding visual processing in the brain is building models that accurately predict neural responses to arbitrary stimuli [1]. Primary visual cortex (V1) has been a strong focus of sensory neuroscience ever since Hubel and Wiesel’s seminal studies demonstrated that neurons in primary visual cortex (V1) respond selectively to distinct image features like local orientation and contrast [2, 3]. Our current standard model of V1 is based on linear-nonlinear models (LN) [4, 5] and energy models [6] to explain simple and complex cells, respectively. While these models work reasonably well to model responses to simple stimuli such as gratings, they fail to account for neural responses to more complex patterns [7] and natural images [8, 9]. Moreover, the computational advantage of orientation-selective LN neurons over simple center-surround filters found in the retina would be unclear [10].

There are a number of hypotheses about nonlinear computations in V1, including normative models like overcomplete sparse coding [11, 12] or canonical computations like divisive normalization [13, 14]. The latter has been used to explain specific phenomena such as center-surround interactions with carefully designed stimuli [15–18]. However, to date, these ideas have not been turned into predictive models of spiking responses that generalize beyond simple stimuli – especially to natural images.

To go beyond simple LN models for natural stimuli, LN-LN cascade models have been proposed, which either learn (convolutional) subunits [19–21] or use handcrafted wavelet representations [22]. These cascade models outperform simple LN models, but they currently do not capture the full range of nonlinearities observed in V1, like gain control mechanisms and potentially other not-yet-understood nonlinear response properties. Because experimental time is limited, LN-LN models have to be designed very carefully to keep the number of parameters tractable, which currently limits their expressiveness, essentially, to energy models for direction-selective and complex cells.

Thus, to make progress in a quantitative sense, recent advances in machine learning and computer vision using deep neural networks (‘deep learning’) have opened a new door by allowing us to learn much more complex nonlinear models of neural responses. There are two main approaches, which we refer to as *goal-driven* and *data-driven*.

The idea behind the goal-driven approach is to train a deep neural network on a high-level task and use the resulting intermediate representations to model neural responses [23, 24]. In the machine learning community, this concept is known as transfer learning and has been very successful in deep learning [25, 26]. Deep convolutional neural networks (CNNs) have reached human-level performance on visual tasks like object classification by training on over one million images [27–30]. These CNNs have proven extremely useful as nonlinear feature spaces for tasks where less labeled data is available [25,31]. This transfer to a new task can be achieved by (linearly) reading out the network’s internal representations of the input. Yamins, DiCarlo and colleagues showed recently that using deep networks trained on large-scale object recognition as nonlinear feature spaces for neural system identification works remarkably well in *higher areas* of the ventral stream, such as V4 and IT [32, 33]. Other groups have used similar approaches for early cortical areas using fMRI [34–36]. However, this approach has not yet been used to model spiking activity of early stages such as V1.

The deep data-driven approach, on the other hand, is based on fitting all model parameters directly to neural data [37–41]. The most critical advance of these models in neural system identification is that they can have many more parameters than the classical LN cascade models discussed above, because they exploit computational similarities between different neurons [38, 40]. While previous approaches treated each neuron as an individual multivariate regression problem, modern CNN-based approaches learn one model for an entire population of neurons, thereby exploiting two key properties of local neural circuits: (1) they share the same presynaptic circuitry (for V1: retina and LGN) [38] and (2) many neurons perform essentially the same computation, but at different locations (topographic organization, implemented by convolutional weight sharing) [39–41].

While both the goal-driven and the data-driven approach have been shown to outperform LN models in some settings, neither approach has been evaluated on spiking activity in monkey V1 (see [42, 43] for concurrent work). In this paper, we fill this gap and evaluate both approaches in monkey V1. We found that deep neural networks lead to substantial performance improvements over older models. In our natural image dataset, goal-driven and data-driven models performed similarly well. The goal-driven approach reached this performance with as little as 20% of the dataset and its performance saturated thereafter. In contrast, the data-driven approach required the full dataset for maximum performance, suggesting that it could benefit from a larger dataset and reach even better performance. Our key finding is that the best models required at least four nonlinear processing steps, suggesting that we need to revise our view of V1 as a Gabor filter bank and appreciate the nonlinear nature of its computations. We conclude that deep networks are not just one among many approaches that can be used, but are – despite their limitations – currently the single most accurate model of V1 computation.

## Results

We measured the spiking activity of populations of neurons in V1 of two awake, fixating rhesus macaques using a linear 32-channel array spanning all cortical layers (Fig 2A). Monkeys were viewing stimuli that consisted of 1450 natural images and four sets of textures synthesized to keep different levels of higher-order correlations present in these images (Fig 1, see Methods). Each trial consisted of a sequence of images shown for 60 ms each, with no blanks in between. In each session, we centered the stimuli on the population receptive field of the neurons.

**Fig 1.**
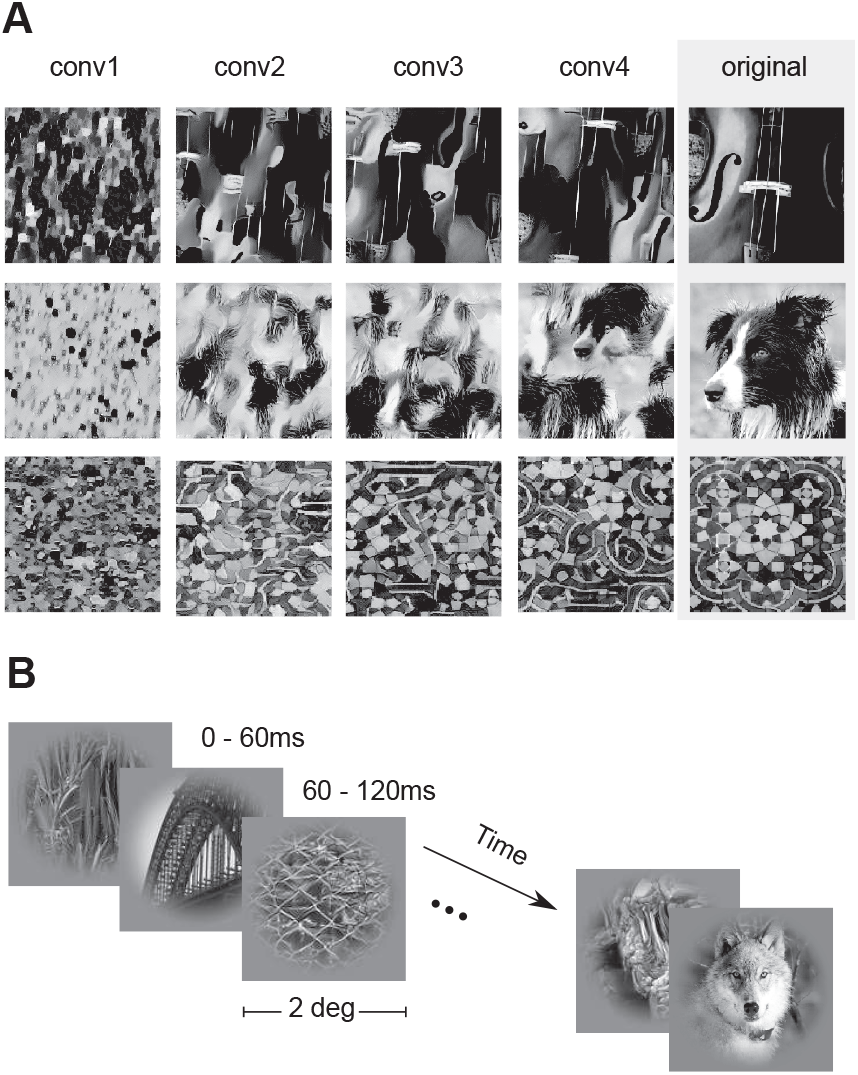
Stimulus paradigm. A: Classes of images shown in the experiment. We used grayscale natural images (labeled ‘original’) from the ImageNet dataset [44] along with textures synthesized from these images using the texture synthesis algorithm described by [45]. Each row shows four synthesized versions of three example original images using different convolutional layers (see Materials and Methods for details). Lower convolutional layers capture more local statistics compared to higher ones. B: Stimulus sequence. In each trial, we showed a randomized sequence of images (each displayed for 60 ms covering 2 degrees of visual angle) centered on the receptive fields of the recorded neurons while the monkey sustained fixation on a target. The images were masked with a circular mask with cosine fadeout.

**Fig 2.**
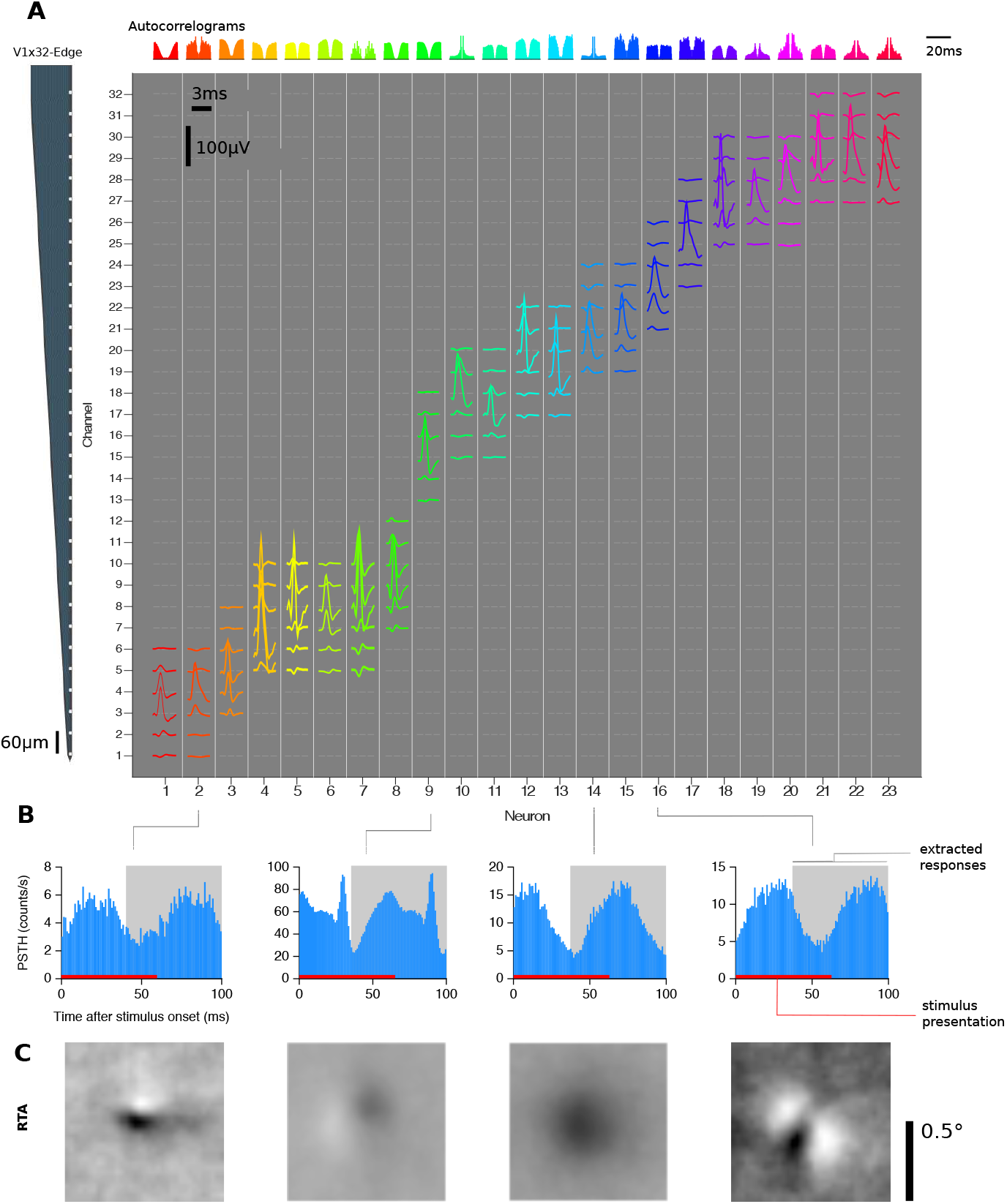
V1 electrophysiological responses. A: Isolated single-unit activity. We performed acute recordings with a 32-channel, linear array (NeuroNexus V1×32-Edge-10mm-60-177, layout shown in the left) to record in primary visual cortex of two awake, fixating macaques. The channel mean-waveform footprints of the spiking activity of 23 well-isolated neurons in one example session are shown in the central larger panel. The upper panel shows color-matched autocorrelograms. B: Peri-stimulus time histograms (PSTH) of four example neurons from A. Spike counts where binned with t = 1 ms, aligned to the onset of each stimulus image, and averaged over trials. The 60 ms interval where the image was displayed is shown in red. We ignored the temporal profile of the response and extracted spike counts for each image on the 40–100 ms interval after image onset (shown in light gray). C: The Response Triggered Average (RTA) calculated by reverse correlation of the extracted responses

We isolated 262 neurons in 17 sessions. The neurons responded well to the fast succession of natural images with a typical response latency of 40ms (Fig. 2B). Therefore, we extracted the spike counts in the window 40-100 ms after image onset (Fig. 2B). The recorded neurons were diverse in their temporal response properties (e.g. see autocorrelogram Fig. 2A), average firing rates in response to stimulus (21.1 ± 20.8 spikes/s, mean ± S.D.), cortical depth (55% of cells in granular, 18% in supragranular, and 27% infragranular layers), and response-triggered average (RTA) structure (Fig. 2C), but neurons recorded on the same array generally had their receptive fields at similar locations approximately centered on the stimulus (Fig. 2C). Prior to analysis, we selected neurons based on how reliable their responses were from trial to trial and included only neurons for which at least 15% of their total variance could be attributed to the stimulus (see Methods). This selection resulted in 166 neurons, which form the basis of the models we describe in the following.

### Generalized linear model with pre-trained CNN features

We start by investigating the goal-driven approach [23, 24]. Here, the idea is to use a high-performing neural network trained on a specific goal – object recognition in this case – as a non-linear feature space and train only a simple linear-nonlinear readout. We chose VGG-19 [28] over other neural networks, because it has a simple architecture (described below), a fine increase in receptive field size along its hierarchy and reasonably high classification accuracy.

VGG-19 is a CNN trained on the large image classification task ImageNet (ILSVRC2012) that takes an RGB image as input and infers the class of the dominant object in the image (among 1000 possible classes). The architecture of VGG-19 consists of a hierarchy of linear-nonlinear transformations (layers), where the input is spatially convolved with a set of filters and then passed through a rectifying nonlinearity (Fig. 3). The output of this operation is again an image with multiple channels. However, these channels do not represent color – as the three channels in the input image – but learned features. They are therefore also called feature maps. Each feature map can be viewed as a filtered version of its input. The collection of such feature maps serves as input for the next layer. Additionally, the network has five pooling layers, where the feature maps are downsampled by a factor of two by taking the local maximum value of four neighboring pixels. There are 16 convolutional layers that can be grouped into five groups named conv1 to conv5 with 2, 2, 4, 4, 4 convolutional layers and 64, 128, 256, 512, 512 output feature maps, respectively, and a pooling layer after each group.

**Fig 3.**
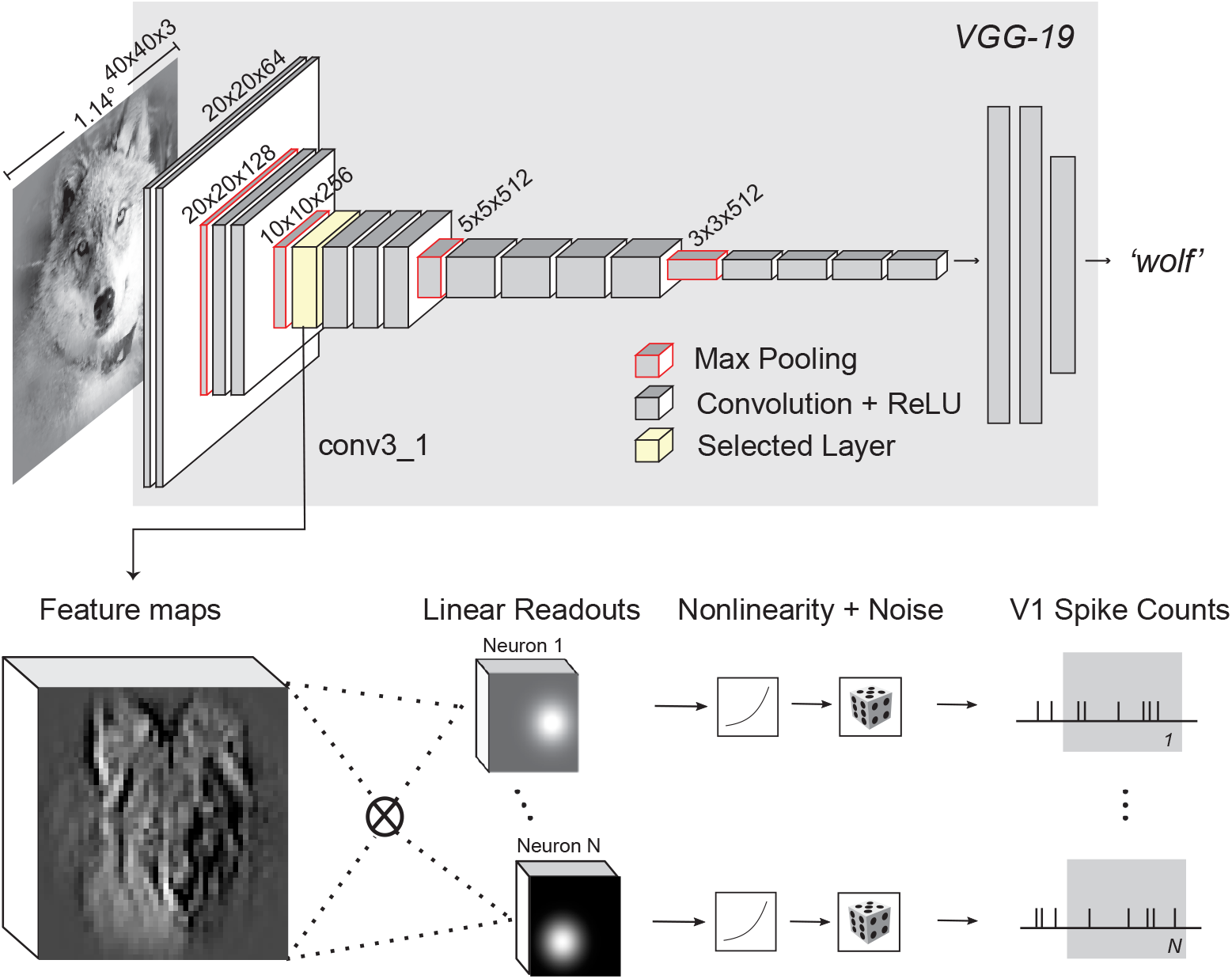
Our proposed model based on VGG-19 features. VGG-19 [28] (gray background) is a trained CNN that takes an input image and produces a class label. For each of the 16 convolutional layers of VGG-19, we extract the feature representations (feature maps) of the images shown to the monkey. We then train for each recorded neuron and convolutional layer, a Generalized Linear Model (GLM) using the feature maps as input to predict the observed spike counts. The GLM is formed by a linear projection (dot product) of the feature maps, a pointwise nonlinearity, and an assumed noise distribution (Poisson) that determines the optimization loss for training. We additionally imposed strong regularization constraints on the readout weights (see text).

We used VGG-19 as a feature space in the following way: We selected the output of a convolutional layer as input features for a Generalized Linear Model (GLM) that predicts the recorded spike counts (Fig. 3). Specifically, we fed each image *x* in our stimulus set through VGG-19 to extract the resulting feature maps *ϕ*(*x*) of a certain layer. These feature maps were then linearly weighted with a set of learned readout weights *w*. This procedure resulted in a single scalar value for each image that was then passed through a (static) output nonlinearity to produce a prediction for the firing rate:

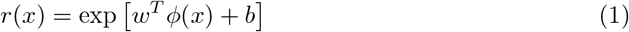

We assumed this prediction to be the mean rate of a Poisson process (see Methods for details). In addition, we applied a number of regularization terms on the readout weights that we explain later.

### Intermediate layers of VGG best predict V1 responses

We first asked which convolutional layer of VGG-19 provides the best feature space for V1. To answer this question, we fitted a readout for each layer and compared the performance. We measured performance by computing the fraction of explainable variance explained (*FEV*). This metric, which ranges from zero to one, measures what fraction of the stimulus-driven response is explained by the model, ignoring the unexplainable trial-to-trial variability in the neurons’ responses (for details see Methods).

We found that the fifth (out of sixteen) layers’ features (called ‘conv3 1’, Fig 3) best predicted neuronal responses to novel images not seen during training (Fig 4, solid line). This model predicted on average 51.6% of the explainable variance. In contrast, performance for the very first layer was poor (31% FEV), but increased monotonically up to conv3 1. Afterwards, the performance again decreased continually up the hierarchy (Fig 4). These results followed our intuition that early to intermediate processing stages in a hierarchical model should match primary visual cortex, given that V1 is the third processing stage in the visual hierarchy after the retina and the lateral geniculate nucleus (LGN) of the thalamus.

**Fig 4.**
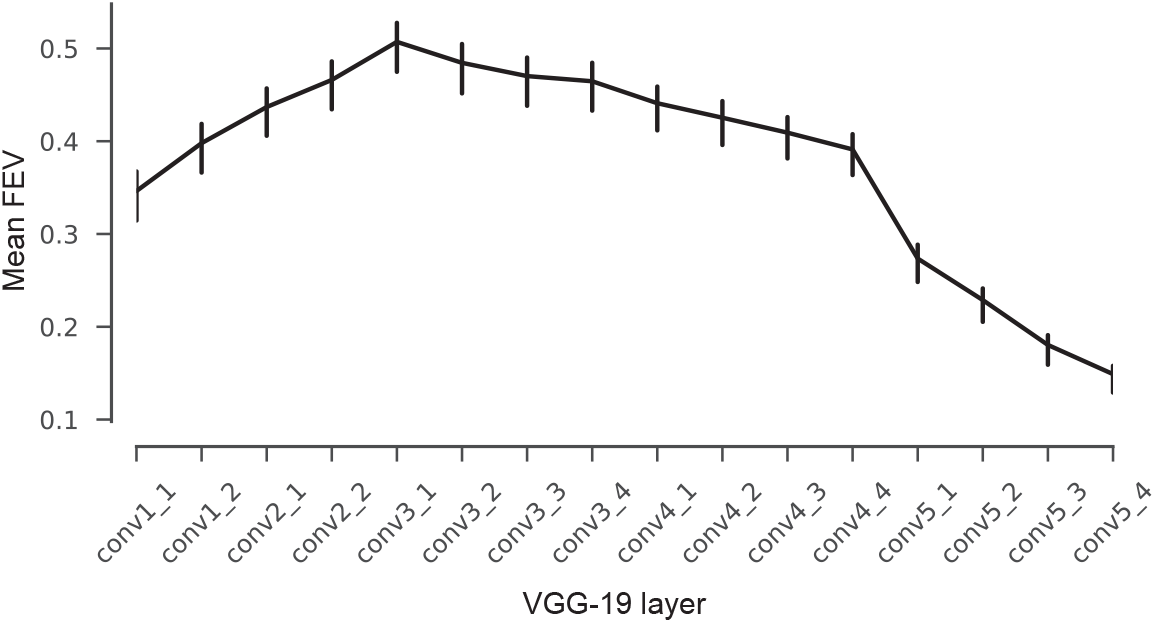
Model performance on test set. Average fraction of explainable variance explained (*FEV*) for models using different VGG layers as nonlinear feature spaces for a GLM. The model based on layer conv3 1 shows on average the highest predictive performance.

### Control for input resolution and receptive field sizes

An important issue to be aware of is that the receptive field sizes of VGG units grow along the hierarchy – just like those of visual neurons in the brain. Incidentally, the receptive fields of units in the best-performing layer conv3 1 subtended approximately 0.68 degrees of visual angle, roughly matching the expected receptive sizes of our V1 neurons given their eccentricities between 1 and 3 degrees. Because receptive fields in VGG are defined in terms of image pixels, their size in degrees of visual angle depends on the resolution at which we present images to VGG, which is a free parameter whose choice will affect the results.

VGG-19 was trained on images of 224×224 px. Given the image resolution we used for the analyses presented above, an entire image would subtend ∼6.4 degrees of visual angle (the crops shown to the monkey were 2 degrees; see Methods for details). Although this choice appears to be reasonable and consistent with earlier work [33], it is to some extent arbitrary. If we had presented the images at lower resolution, the receptive fields sizes of all VGG units would have been larger. As a consequence, the receptive fields of units in earlier layers would match those of V1 and these layers may perform better. If this was indeed the case, there would be nothing special about layer conv3 1 with respect to V1.

To ensure that the choice of input resolution did not affect our results, we performed a control experiment, which substantiated our claim that con3 1 provides the best features for V1. We repeated the model comparison presented above with different input resolutions, rescaling the image crops by a factor of 0.67 and 1.5. These resolutions correspond to 9.55 and 4.25 degrees of full visual field for VGG-19, respectively. While changing the input resolution did shift the optimal layer towards that with matching receptive field sizes (Fig. 5, first and third row), the resolution we had picked for our main experiment yielded the best overall performance (Fig. 5, second row, third column). Thus, over a range of input resolutions and layers, conv3 1 performed best, although conv2 2 at lower resolution yielded only slightly lower performance.

**Fig 5.**
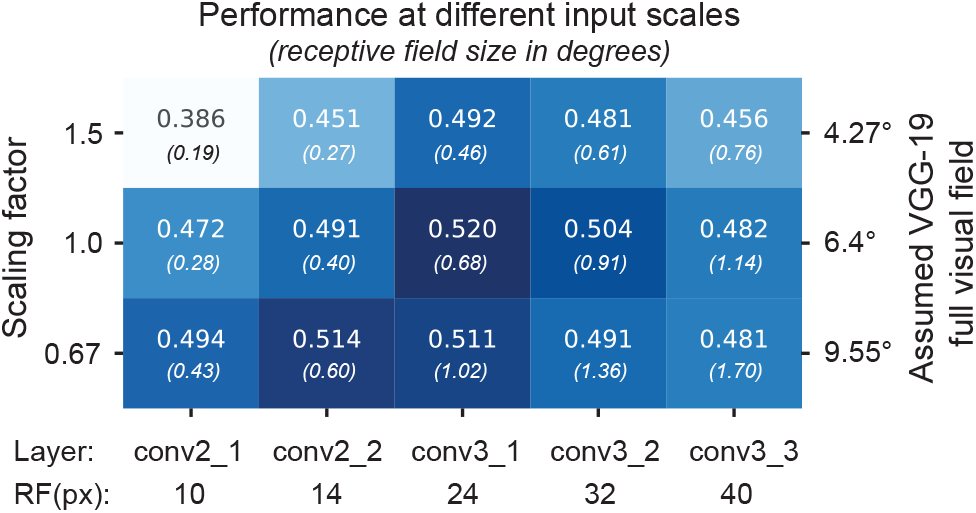
VGG-19 based model performance at different input scales. The performance on test set of cross-validated models that use as feature spaces layers conv2 1 to conv3 3 for different input resolutions. With the original scale used in Fig. 4, we assumed that VGG-19 was trained with 6.4 degrees field of view. 0.67 and 1.5 times the original scale justify the choice of resolution for further analysis. At the bottom, the receptive field sizes of the different layers.

### Careful regularization is necessary

The number of predictors given by the convolutional feature space of a large pre-trained network is much larger than the number of pixels in the image. Most of these predictors will likely be irrelevant for most recorded neuron –– for example, network units at spatial positions that are not aligned with the neuron’s receptive field or feature maps that compute nonlinearities unrelated to those of the cells. Naïvely including many unimportant predictors would prevent us from learning a good mapping, because they lead to overfitting. We therefore used a regularization scheme with the following three terms for the readout weights: (1) sparsity, to encourage the selection of a few units; (2) smoothness, for a regular spatial continuity of the predictors’ receptive fields; and (3) group sparsity, to encourage the model to pool from a small number of feature maps (see Methods for details).

We found that regularization was key to obtaining good performance (Table 1). The full model with all three terms had the best performance on the test set and vastly outperformed a model with no regularization. Eliminating one of the three terms while keeping the other two hurt performance only marginally. Among the three regularizers, sparsity appeared to be the most important one quantitatively, whereas smoothness and group sparsity could be dropped without hurting overall performance.

**Table 1.** Ablation experiments for VGG-based model, removing regularization terms (rows 2–5) and using factorized readout weights (row 6, [40]).

| Model | FEV |
| --- | --- |
| **Full model** | **0.52** |
| No smoothness | 0.51 |
| No sparsity | 0.49 |
| No group sparsity | 0.51 |
| No regularization | 0.33 |
| Factorized readout [40] | 0.45 |

To understand the effect of the different regularizers qualitatively, we visualized the readout weights of each feature map of our conv3 1-based model, ordered by their spatial energy for each cell, for each of the regularization schemes (see Fig. 6A for five sample neurons). Without the sparsity constraint, we obtained smooth but spread-out weights that were not well localized. Dropping the smoothness term – despite performing equally in a quantitative sense – produced sparse activations that were less localized and not smooth. Without any regularization, the weights appeared noisy and one could not get any insights about the locality of the neuron. On the other hand, the full model –– in addition to having the best performance –– also provides localized and smooth filters that provide information about the neurons’ receptive field and the set of useful feature maps for prediction.

**Fig 6.**
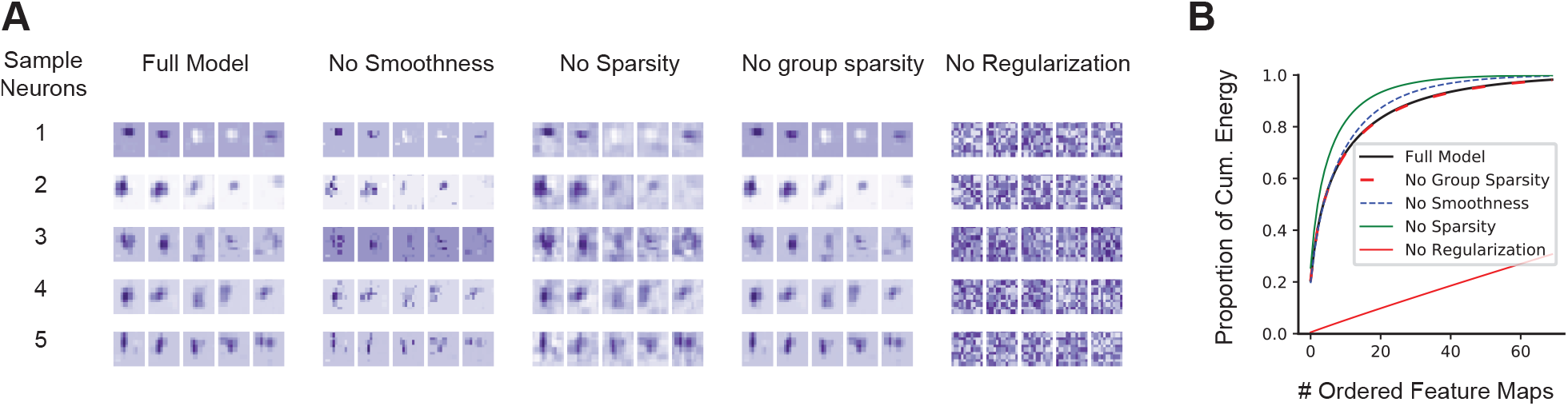
Learned readout weights with different regularization modes. **A**. For five example neurons (rows), the five highest-energy spatial readouts out of 256 feature maps of conv3 1 for each regularization scheme we explored. The full model exhibits the most localized and smooth receptive fields. **B**. The highest normalized spatial energy of the learned readouts as a function of ordered feature maps (first 70 out of 256 of conv3 1 shown) averaged for all cells. With regularization, only a few feature maps are used for prediction, quickly asymptoting at 1.

Finally, we also observed that only a small number of feature maps was used for each neuron: the weights decayed exponentially and only 20 feature maps out of 256 contained on average 82% of the readout energy (Fig. 6B).

An alternative form of regularization or inductive bias would be to constrain the readout weights to be factorized in space and features [40], which reduces the number of parameters substantially. However, the best model with this factorized readout achieved only 45.5% FEV (Table 1), presumably because the feature space has not been optimized for such a constrained readout.

### Goal-driven and data driven CNNs set the state of the art

Multi-layer feedforward networks have been fitted successfully to neural data on natural image datasets in mouse V1 [38, 40]. Thus, we inquired how our goal-driven model compares to a model belonging to the same functional class, but directly fitted to the neural data. Following the methods proposed by Klindt et. al [40], we fitted CNNs with one to five convolutional layers (Fig 7A; see Methods for details).

**Fig 7.**
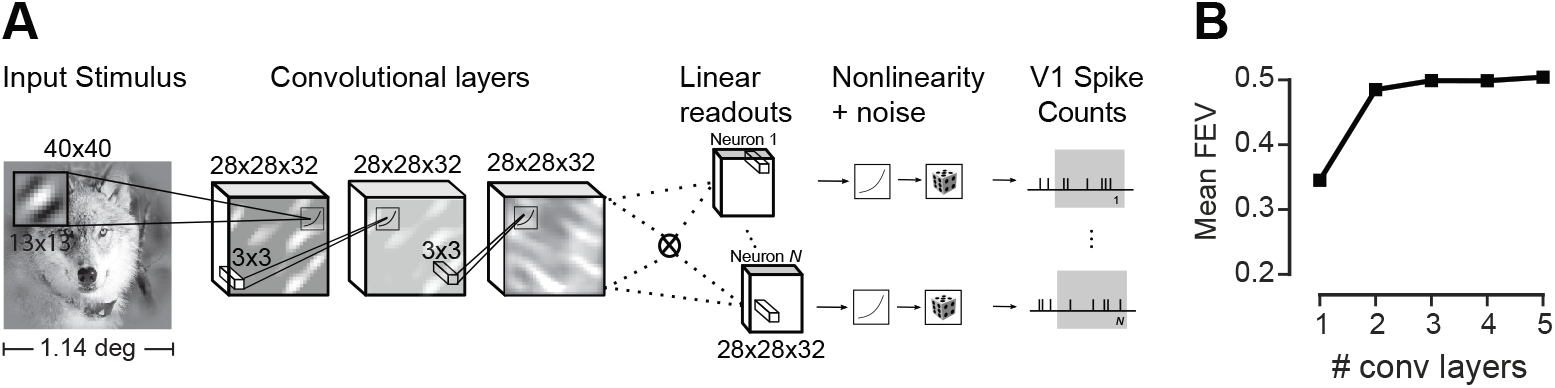
Data-driven convolutional network model. We trained a convolutional neural network to produce a feature space fed to a GLM-like model. In contrast to the VGG-based model, both feature space and readout weights are trained only on the neural data. **A**. Three-layer architecture with a factorized readout [40] used for comparison with other models. **B**. Performance of the data driven approach as a function of the number of convolutional layers on held-out data. Three convolutional layers provided the best performance on the validation set. See methods for details.

The data-driven CNNs with three or more convolutional layers yielded the best performance, outperforming their competitors with fewer (one or two) layers (Fig 7B). We therefore decided to use the CNN with three layers for model comparison, as it is the simplest model with highest predictive power on the validation set.

We then asked how the predictive performance of both data-driven and goal driven models compares to previous models of V1. As a baseline, we fitted a regularized version of the classical linear-nonlinear Poisson model (LNP; [46]). The LNP is a very popular model used to estimate the receptive field of neurons and offers interpretability and convexity for its optimization. This model gave us a good idea of the nonlinearity of the cells’ responses. Additionally, we fit a model based on a handcrafted nonlinear feature space consisting of a set of Gabor wavelets [4, 47–49] and energy terms of each quadrature pair [6]. We refer to this model as the ‘Gabor filter bank’ (GFB). It builds upon existing knowledge about V1 function and is able to model simple and complex cells. Moreover, this model is the current state of the art in the neural prediction challenge for monkey V1 responses to natural images [50] and therefore a strong baseline for a quantitative evaluation.

We compared the models for a number of cells from a representative recording (Fig 8A). There was a diversity of cells, both in terms of how much variance could be explained in principle (dark gray bars) and how well the individual models performed (colored bars). Overall, the deep learning models consistently outperformed the two simpler models of V1. This trend was consistent across the entire dataset (Fig 8B, D). The LNP model achieved 17% *FEV*, the GFB model 40.2% *FEV*. The performance of the CNN trained directly on the data was comparable to that of the VGG-based model (Fig 8C, D); they predicted 50.0% and 51.5% *FEV*, respectively, on average. Note that the one-layer CNN (mean 34.5% *FEV*, Fig. 7) structurally resembles the convolutional subunit model proposed by Vintch and colleagues [21]. Thus, deeper CNNs also outperform learned LN-LN cascade models significantly.

**Fig 8.**
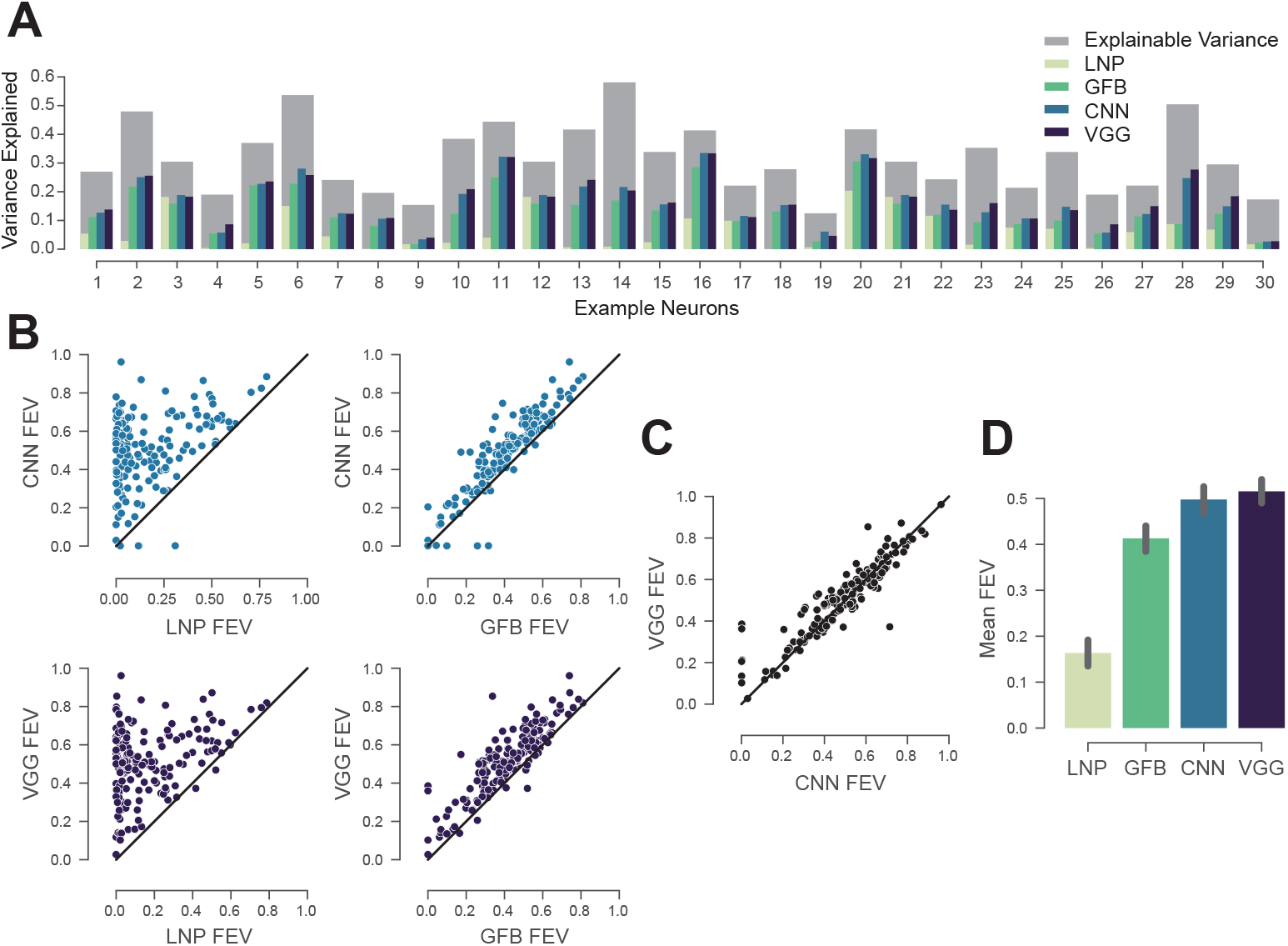
Deep models are the new state of the art. A: Randomly selected cells. The normalized explainable variance (oracle) per cell is shown in gray. For each cell from left to right, the variance explained of: regularized LNP [46], GFB [22, 47, 48], three-layer CNN trained on neural responses, and VGG conv3 1 model (ours). B. CNN and VGG conv3 1 models outperform for most cells LNP and GFB. Black line denotes the identity. The performance is given in *FEV*. C: VGG conv3 1 features perform slightly better than the three-layer CNN. D: Average performance of the four models given in mean fraction of explainable variance explained (*FEV*).

### Improvement of model predictions is not linked to neurons’ tuning properties

We next asked whether the improvement in predictive performance afforded by our deep neural network models was related in any way to known tuning properties of V1 neurons such as the shape of their orientation tuning curve or their classification along the simple-complex axis. To investigate this question, we performed an in-silico experiment: we showed Gabor patches of the same size as our image stimulus with various orientations, spatial frequencies and phases (Fig. 9A) to our CNN model of each cell. Based on the model output, we computed tuning curves for orientation (Fig. 9B) and spatial phase (Fig. 9D) by using the set of Gabors with the optimal spatial frequency for each neuron.

**Fig 9.**
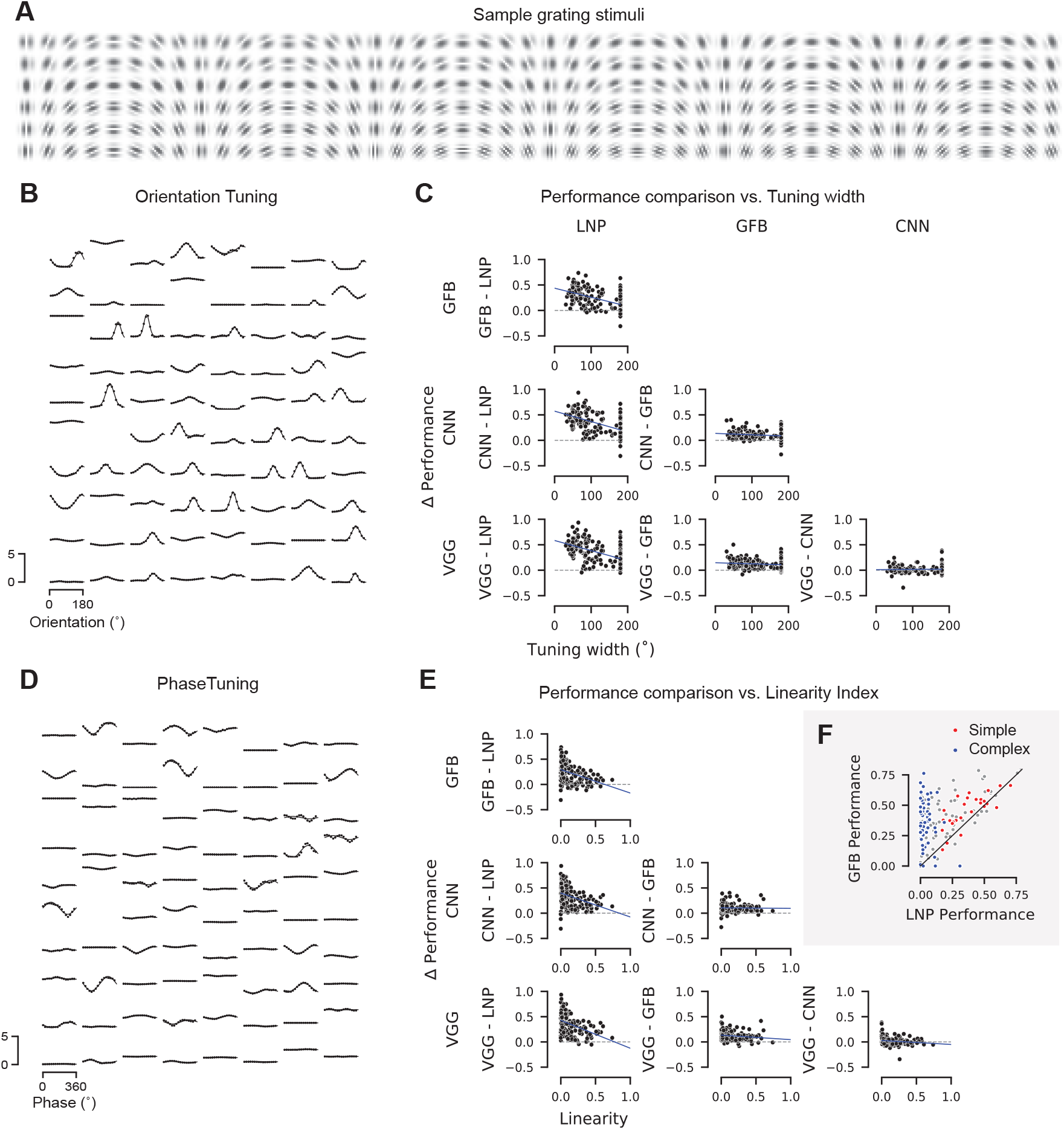
Relationship between model performance and neurons’ tuning properties. **A**. A sample subset of the Gabor stimuli with a rich diversity of frequencies, orientations, and phases. **B**. Dots: Orientation tuning curves of 80 sample neurons predicted by our CNN model. Tuning curves computed at the optimal spatial frequency and phase for each neuron. Lines: von Mises fits (see Methods). **C**. Difference in performance between pairs of the four models as a function of tuning width. Tuning width defined as the full width at half maximum of the fitted tuning curve. **D**. Dots: Phase tuning curves of the same 80 sample neurons as in B, predicted by our CNN model. Tuning curves computed at the optimal spatial frequency and orientation for each neuron. Lines: Cosine tuning curve with fitted amplitude and offset (see Methods). **E**. Like C, the difference in performance between pairs of models as a function of the neurons’ linearity index. Linearity index: ratio of amplitude of cosine over offset (0: complex; 1: simple). **F**. Performance comparison between GFB and LNP model. Red: simple cells (top 16% linearity, linearity > 0.3); blue: complex cells (bottom 28% linearity, linearity *<* 0.04).

Based on the phase tuning curves we compute a linearity index (see Methods), which locates each cell on the axis from simple (linearity index close to one) to complex (index close to zero). We then asked whether there are systematic differences in model performance as a function of this simple-complex characterization. As expected, we found that more complex cells are explained better by the Gabor filter bank model than an LNP model (Fig. 9C). The same was true for both the data-driven CNN and the VGG-based model. However, the simple-complex axis did not predict whether and how much the CNN models outperformed the Gabor filter bank model. Thus, whatever aspect of V1 computation was additionally explained by the CNN models, it was shared by both simple and complex cells.

Next, we asked whether there is a relationship between orientation selectivity (tuning width) and the performance of any of our models. We found that for cells with sharper orientation tuning, the performance gain afforded by the Gabor filter bank model (and both CNN-based models) over an LNP was larger than for less sharply tuned cells (Fig. 9E). This result is not unexpected given that cells in layer 2/3 tend to have narrower tuning curves and also tend to be more complex (refs). However, as for the simple-complex axis, tuning width was not predictive of the performance gain afforded by a CNN-based model over the Gabor filter bank (Fig. 9E). Therefore, any additional nonlinearity in V1 computation captured by the CNN models is not specific to sharply or broadly tuned neurons.

### Models generalize across stimulus statistics

Our stimulus set contains both natural images as well as four sets of textures generated from those images. These textures differ in how accurately and over what spatial extent they reproduce the local image statistics (see Fig. 1). On the one end of the spectrum, samples from the conv1 model reproduce relatively linear statistics over small regions of a few minutes of arc. On the other end of the spectrum, samples from the conv4 model almost perfectly reproduce the statistics of natural images over larger regions of 1–2 degrees of visual angle, covering the entire classical and at least part of the extra-classical receptive field of V1 neurons.

We asked to what extent including these different image statistics helps or hurts building a predictive model. To answer this question, we additionally fit both the data-driven CNN model and the VGG-based model to subsets of the data containing only images from a single image type (originals or one of four texture classes). We then evaluated each of these models on all image types (Fig. 10). Perhaps surprisingly, we found that using any of the four texture statistics or the original images for training lead to approximately equal performance, independent on which images were used for testing the model (Fig. 10). This result held for both the VGG-based (Fig. 10A) and the data-driven CNN model (Fig. 10B). Thus, using the very localized conv1 textures worked just as well for predicting the responses to natural images as did training directly on natural images – or any other combination of training and test set. This result is somewhat surprising to us, as the conv1 textures match only very simple and local statistics on spatial scales smaller than individidual neurons’ receptive fields and perceptually are much closer to noise than natural images.

**Fig 10.**
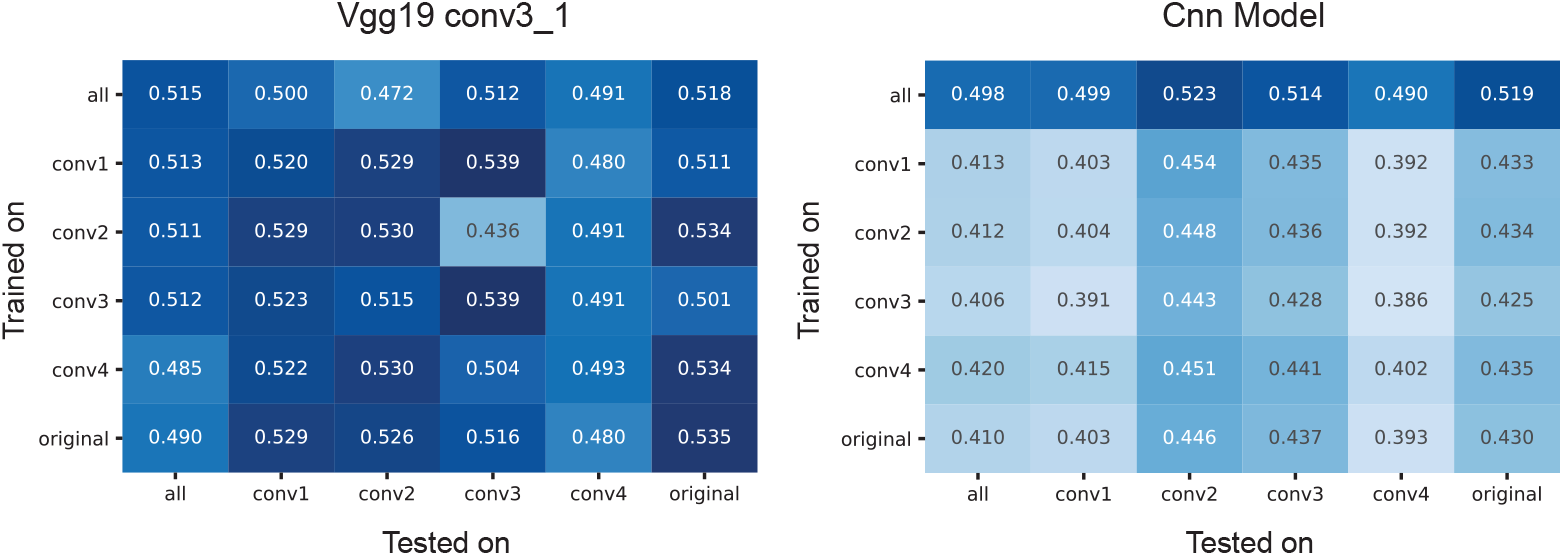
Training and evaluation on the different stimulus types. For both conv3 1 features of VGG-19 (left) and CNN-based models, we trained with all and every individual stimulus type (rows) (see Fig 1) and tested on all and every individual type. The VGG model showed good domain transfer in general. The same was true for the data-driven CNN model, although it performed worse overall when trained on only one set of images due to the smaller training sample. There were no substantial differences in performance across image statistics.

### VGG-based model needs less training data

An interesting corollary of the analysis above is the difference in absolute performance between the VGG-based and the data-driven CNN model when using only a subset of images for training: while the performance of the VGG-based model remains equally high when using only a fifth of the data for training (Fig. 10A), the data-driven CNN takes a substantial hit (Fig. 10B, second and following rows). Thus, while the two models perform similarly when using our entire dataset, the VGG-based model works better when less training data is available. This result indicates that for our current experimental paradigm training the readout weights is not the bottleneck – despite the readout containing a large number of parameters in the VGG-based model (Table 2). Because we know that only a small number of non-zero weights are necessary, the L1 regularizer works very well in this case. In contrast, the data-driven model takes a substantial hit when using only a subset of the data, suggesting that learning the shared feature space is the bottleneck for this model. Thus collecting a larger dataset could help the data-driven model but is unlikely to improve performance of the VGG-based one.

**Table 2.**
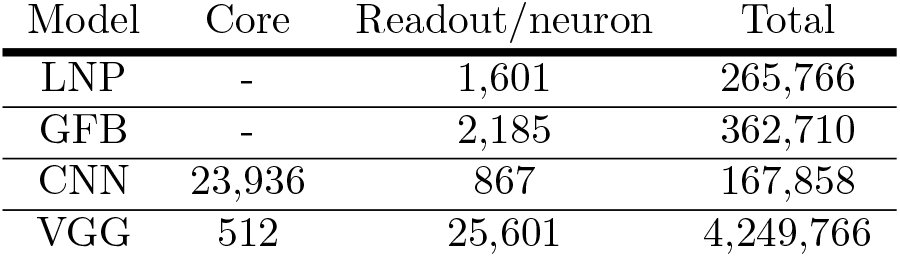
Number of learned parameters for the different models. ‘Core’ refers to the part shared among all neurons. ‘Readout’ refers to the parameters required for each neuron.

| Model | Core | Readout/neuron | Total |
| --- | --- | --- | --- |
| LNP | - | 1,601 | 265,766 |
| GFB | - | 2,185 | 362,710 |
| CNN | 23,936 | 867 | 167,858 |
| VGG | 512 | 25,601 | 4,249,766 |

## Discussion

Our goal was to find which model among various alternatives is best for one of the most studied systems in modern systems neuroscience: primary visual cortex. We fit two models based on convolutional neural networks to V1 responses to natural stimuli in awake, fixating monkeys: a goal-driven model, which uses the representations learned by a CNN trained on object recognition (VGG-19), and a data-driven model, which learns both the convolutional and readout parameters using stimulus-response pairs with multiple neurons simultaneously. Both approaches yielded comparable performance and substantially outperformed the widely used LNP [46] and a rich Gabor filter bank (GFB), which held the previous state of the art in prediction of V1 responses to natural images. This finding is of great importance because it suggests that deep neural networks can be used to model not only higher cortex, but also lower cortical areas. In fact, deep networks are not just one among many approaches that can be used, but the only class of models that has been shown to provide the multiple nonlinearities necessary to accurately describe V1 responses to natural stimuli.

Our work contributes to a growing body of research where goal-driven deep learning models [23,24] have shown unprecedented predictive performance in higher areas of the visual stream [32, 33], and a hierarchical correspondence between deep networks and the ventral stream [35, 51]. Studies based on fMRI have established a correspondence between early layers of CNNs trained on object recognition and V1 [35, 52]. Here, with electrophysiological data and a deeper network (VGG-19), we found that V1 is better explained by feature spaces multiple nonlinearities away from the pixels. We found that it takes five layers (a quarter of the way) into the computational stack of the categorization network to explain V1 best, which is in contrast to the many models that treat V1 as only one or two nonlinearities away from pixels (i.e. GLMs, energy models). Earlier layers of our CNNs might explain subcortical areas better (i.e. retina and LGN), as they are known to be modeled best with multiple, but fewer, nonlinearities already [41].

What are, then, the additional nonlinearities captured by our deep convolutional models beyond those in LNP or GFB? Our first attempts to answer this question via an in-silico analysis revealed that whatever the CNNs capture beyond the Gabor filter bank model is not specific to the cells’ tuning properties, such as width of the orientation tuning curve and their characterization along the simple-complex spectrum. This result suggests that the missing nonlinearity may be relatively generic and applicable to most cells. There are a few clear candidates for such nonlinear computations, including divisive normalization [53] and overcomplete sparse coding [12]. Unfortunately, quantifying whether these theories provide an equally good account of the data is not straightforward: so far they have not been turned into predictive models for V1 neurons that are applicable to natural images. In the case of divisive normalization, the main challenge is learning the normalization pool. There is evidence for multiple normalization pools, both tuned and untuned and operating in the receptive field center and surround [54]. However, previous work investigating these normalization pools has employed simple stimuli such as gratings [18] and we are not aware of any work learning the entire normalization function from neural responses to natural stimuli. Similarly, sparse coding has so far been evaluated only qualitatively by showing that the learned basis functions resemble Gabor filters [12]. Solving a convolutional sparse coding problem [55] and using the resulting representation as a feature space would be a promising direction for future work, but we consider re-implementing and thoroughly evaluating this approach to be beyond the scope of the current paper.

To move forward in understanding such nonlinearities may require developing more interpretable neural networks or methods that provide interpretability of networks, which are an active area of research in the machine learning community. Alternatively, we could build predictive models constrained with specific hard-coded nonlinearities (such as normalization) that express our knowledge about important computations.

It is also possible that the mechanistic level of circuit components remains underconstrained by function and thus allows only for explanations up to some degree of degeneracy, requiring knowledge of the objective function the system optimizes (e.g. sparse coding, predictive coding). Our results show that object categorization – despite being a relatively impoverished visual task – is a very useful learning objective not only for high-level areas in the ventral stream, but also for a more low-level and general-purpose area like V1, despite the fact that V1 clearly serves a large number of tasks beyond object categorization. This finding resonates well with results from computer vision, where object categorization has also been found to be an extremely useful objective to learn features applicable to numerous other visual tasks [25].

Our current best models still leave almost half of the explainable variance unexplained, raising the question of how to make further progress. Our finding that the VGG-based model performed equally well with only 20% of the images in the training set suggests that its performance was not limited by the amount of data available to learn the readout weights, which make for the bulk of the parameters in this model (Table 2). Instead, the VGG-based model appears to be limited by a remaining mismatch between VGG features and V1 computation. This mismatch could potentially be reduced by using features from neural networks trained simultaneously on multiple ethologically relevant tasks beyond object categorization. The data-driven model reached its full performance only with the full training set, suggesting that learning the nonlinear feature space is the bottleneck. In this case, pooling over a larger number of neurons or recording longer from the same neurons should improve performance because most of the parameters are in the shared feature space (Table 2) and this number is independent of the number of neurons being modeled.

We conclude that previous attempts to describe the basic computations that different types of neurons in primary visual cortex perform (e.g.“edge detection”) do not account for the complexity of multi-layer nonlinear computations that are necessary for the performance boost achieved with CNNs. Although these models, which so far best describe these computations, are complex and lack a concise intuitive description, they can be obtained by a simple principle: optimize a network to solve an ecologically relevant task (object categorization) and use the hidden representations of such a network. For future work, combining data- and goal-driven models and incorporating the recurrent lateral and feedback connections of the neocortex promise to provide a framework for incrementally unravelling the nonlinear computations of V1 neurons.

## Methods

### Electrophysiological recordings

All behavioral and electrophysiological data were obtained from two healthy, male rhesus macaque (Macaca mulatta) monkeys aged 12 and 9 years and weighing 12 and 10 kg, respectively, during the time of study. All experimental procedures complied with guidelines of the NIH and were approved by the Baylor College of Medicine Institutional Animal Care and Use Committee (permit number: AN-4367). Animals were housed individually in a large room located adjacent to the training facility, along with around ten other monkeys permitting rich visual, olfactory and auditory interactions, on a 12h light/dark cycle. Regular veterinary care and monitoring, balanced nutrition and environmental enrichment were provided by the Center for Comparative Medicine of Baylor College of Medicine. Surgical procedures on monkeys were conducted under general anesthesia following standard aseptic techniques. To ameliorate pain after surgery, analgesics were given for 7 days. Animals were not sacrificed after the experiments.

We performed non-chronic recordings using a 32-channel linear silicon probe (NeuroNexus V1×32-Edge-10mm-60-177). The surgical methods and recording protocol were described previously [56]. Briefly, form-specific titanium recording chambers and headposts were implanted under full anesthesia and aseptic conditions. The bone was originally left intact and only prior to recordings, small trephinations (2 mm) were made over medial primary visual cortex at eccentricities ranging from 1.4 to 3.0 degrees of visual angle. Recordings were done within two weeks of each trephination. Probes were lowered using a Narishige Microdrive (MO-97) and a guide tube to penetrate the dura. Care was taken to lower the probe slowly, not to penetrate the cortex with the guide tube and to minimize tissue compression (for a detailed description of the procedure, see [56]).

### Data acquisition and spike sorting

Electrophysiological data were collected continuously as broadband signal (0.5Hz–16kHz) digitized at 24 bits as described previously [57]. Our spike sorting methods are based on [58], code available at https://github.com/aecker/moksm, but with adaptations to the novel type of silicon probe as described previously [56]. Briefly, we split the linear array of 32 channels into 14 groups of 6 adjacent channels (with a stride of two), which we treated as virtual electrodes for spike detection and sorting. Spikes were detected when channel signals crossed a threshold of five times the standard deviation of the noise. After spike alignment, we extracted the first three principal components of each channel, resulting in an 18-dimensional feature space used for spike sorting. We fitted a Kalman filter mixture model [59, 60] to track waveform drift typical for non-chronic recordings. The shape of each cluster was modeled with a multivariate *t*-distribution (*df* = 5) with a ridge-regularized covariance matrix. The number of clusters was determined based on a penalized average likelihood with a constant cost per additional cluster [58]. Subsequently, we used a custom graphical user interface to manually verify single-unit isolation by assessing the stability of the units (based on drifts and health of the cells throughout the session), identifying a refractory period, and inspecting the scatter plots of the pairs of channel principal components.

### Visual stimulation and eye tracking

Visual stimuli were rendered by a dedicated graphics workstation and displayed on a 19” CRT monitor (40×30 cm) with a refresh rate of 100 Hz at a resolution of 1600×1200 pixels and a viewing distance of 100 cm (resulting in ~70 px/deg). The monitors were gamma-corrected to have a linear luminance response profile. A camera-based, custom-built eye tracking system verified that monkeys maintained fixation within ∼0.42 degrees around the target. Offline analysis showed that monkeys typically fixated much more accurately. The monkeys were trained to fixate on a red target of ∼0.15 degrees in the middle of the screen. After they maintained fixation for 300 ms, a visual stimulus appeared. If the monkeys fixated throughout the entire stimulus period, they received a drop of juice at the end of the trial.

### Receptive field mapping

At the beginning of each session, we first mapped receptive fields. We used a sparse random dot stimulus for receptive field mapping. A single dot of size 0.12 degrees of visual field was presented on a uniform gray background, changing location and color (black or white) randomly every 30 ms. Each trial lasted for two seconds. We obtained multi-unit receptive field profiles for every channel using reverse correlation. We then estimated the population receptive field location by fitting a 2D Gaussian to the spike-triggered average across channels at the time lag that maximizes the signal-to-noise-ratio. We subsequently placed our natural image stimulus at this location.

### Natural image stimulus

We used a set of 1540 grayscale images as well as four texturized versions of each image. We used grayscale images to avoid the complexity of dealing with color and focus on spatial image statistics. The texturized stimuli allowed us to vary the degree of naturalness, ranging from relatively simple, local statistics to very realistic textures capturing image statistics over spatial scales covering both classical and at least parts of the extra-classical receptive field of neurons. The images were taken from ImageNet [44], converted to grayscale and rescaled to 256×256 pixels. We generated textures with different degrees of naturalness by capturing different levels of higher-order correlations from a local to a global scale by using a parametric model for texture synthesis [45]. This texture model uses summary statistics of feature activations in different layers of the VGG-19 network [28] as parameters for the texture. The lowest-level model uses only the statistics of layer conv1 1. We refer to it as the “conv1” model. The next one uses statistics of conv1 1 and conv2 1 (referred to as conv2), and so on for conv3 and conv4. Due to the increasing level of nonlinearity of the VGG-19 features and their increasing receptive field sizes with depth, the textures synthesized from these models become increasingly more natural (see Fig. 1 and [45] for more examples)

To synthesize the textures, we start with a random white noise image and iteratively refine pixels via gradient descent such that the resulting image matches the feature statistics of the original image [45]. For displaying and further analyses, we cropped the central 140 pixels of each image, which corresponds to 2 degrees of visual angle.

The entire data set contains 1450×5 = 7250 images (original plus synthesized). During each trial, 29 images were displayed, each for 60 ms, with no blanks in between (Fig 1B). We chose this fast succession of images to maximize the number of images we can get through in a single experiment, resulting in a large training set for model fitting. The short presentation times also mean that the responses we observe are mainly feedforward, since feedback processes take some time to be engaged. Each image was masked by a circular mask with a diameter of 2 degrees (140 px) and a soft fade-out starting at a diameter of 1 degree:

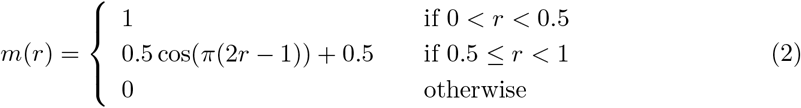

Images were randomized such that consecutive images were not of the same type or synthesized from the same image. A full pass through the dataset took 250 successful trials, after which it was traversed again in a new random order. Images were repeated between one and four times, depending on how many trials the monkeys completed in each session.

### Dataset and inclusion criteria

We recorded a total of 307 neurons in 23 recording sessions. We did not consider six of these sessions, for which we did not obtain enough trials to have at least two repetitions for each image. In the remaining 17 sessions, we quantified the fraction of total variance of each neuron attributable to the stimulus by computing the ratio of explainable and total variance:

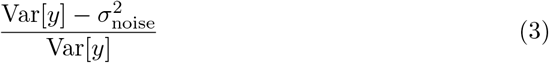

The explainable variance is the total variance minus the variance of the observation noise. We estimated the variance of the observation noise, 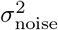, by averaging (across images) the variance (across repetitions) of responses to the same stimulus:

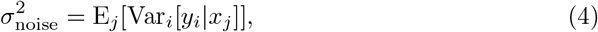

where *x_j_* is the *j*^th^ image and *y_i_* the response to the *i*^th^ repetition. We discarded neurons with a ratio of explainable-to-total variance smaller than 0.15, yielding 166 isolated neurons (monkey A: 51, monkey B: 115) recorded in 17 sessions. Monkey A had only sessions with two repetitions while Monkey B had four repetitions per image.

### Image preprocessing

Before displaying the images on the screen, we normalized them by subtracting the mean intensity of the aperture (unmasked part) across all images and pixels and dividing by its standard deviation (also taken across images and pixels). Then, we scaled the images back with the original full standard deviation and added 128 so that they are between 0 and 255. Prior to model fitting, we additionally cropped the central 80 pixels (1.1°) of the 140-pixel (2°) images shown to the monkey. For most of the analyses presented in this paper, we sub-sampled these crops to half their size (40×40) and z-scored them. For the input resolution control (Fig 5), we resampled with bicubic interpolation the original 80×80 crops to 60×60, 40×40, and 27×27 for scales 1.5, 1, 0.67, respectively.

### GLM with pre-trained CNN features

Our proposed model consists of two parts: feature extraction and a generalized linear model (GLM; Fig 3). The features are the output maps *ϕ*(*x*) of convolutional layers of VGG-19 [28] to a stimulus image *x*, followed by a batch normalization layer. We perform this normalization to ensure that the activations of each feature map have zero mean and unit variance (before ReLU), which is important because the readout weights are regularized by an *L*_1_ penalty and having input features with different variances would implicitly apply different penalties on their corresponding readout weights.

We fit a separate GLM for each convolutional layer of VGG-19. The GLM consists of linear fully connected weights *w_ijk_* for each neuron that compute a dot product with the input feature maps *ϕ_ijk_*(*x*), a static output nonlinearity *f* (also known as the inverse of the link function), and a Poisson noise model used for training. Here, *i* and *j* index space, while *k* indexes feature maps (denoted as depth in Fig 3). The spiking rate of a given neuron *r* will follow:

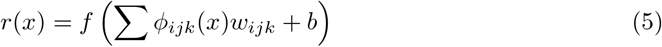

Additionally, three regularization terms were applied to the weights:

1. **Sparsity**: Most weights need to be zero since we expect the spatial pooling to be localized. We use the *L*_1_ norm of the weights:

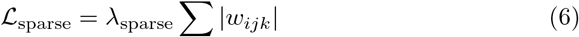
2. **Spatial Smoothness**: Together with sparseness, spatial smoothness encourages spatial locality by imposing continual regular changes in space. We computed this by an *L*_2_ penalty on the Laplacian of the weights:

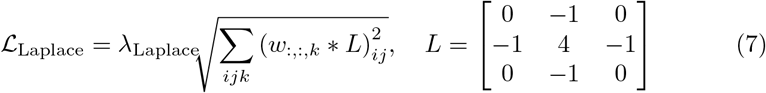
3. **Group Sparsity** encourages our model to pool from a small set of feature maps to explain each neuron’s responses:

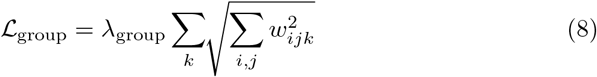

Considering the recorded image-response pair (*x, y*) for one neuron, the resulting loss function is given by:

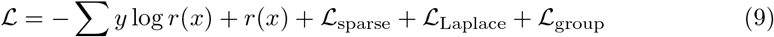

where the sum runs over samples (image, response pairs).

We fit the model by minimizing the loss using the Adam optimizer [61] on a training set consisting of 80% of the data, and reported performance on the remaining 20%. We cross-validated the hyperparameters *λ*_sparse_, *λ*_Laplace_, *λ*_group_ for each neuron independently by performing a grid search over four logarithmically spaced values for each hyperparameter. The validation was done on 20% of the training data. The optimal hyperparameter values obtained on the validation set where *λ*_Laplace_ = 0.1, *λ*_sparse_ = 0.01, *λ_group_* = 0.001. When fitting models, we used the same split of data for training, validation, and testing across all models.

### Data-driven convolutional neural network model

We followed the results of [40] and use their best-performing architecture that obtained state-of-the-art performance on a public dataset [38]. Like our VGG-based model, this model also consisted of convolutional feature extraction followed by a GLM, the difference being that here the convolutional feature space was learned from neural data instead of having been trained on object recognition. The feature extraction architecture consisted of convolutional layers with filters of size 13×13 px for the first layer and 3×3 px for the subsequent layers. Each layer had 32 feature maps (Fig 7A) and exponential linear units (ELU [62])

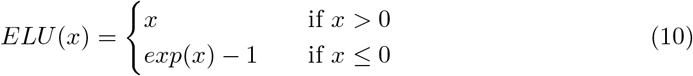

as nonlinearities with batch normalization [63] to facilitate training in between the layers. As in the original publication [40], we regularized the convolutional filters by imposing smoothness constraints on the first layer and group sparseness on the subsequent layers. A notable difference to our VGG-based GLM is that here the readout weights are factorized in space and feature maps:

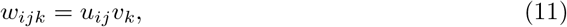

where *u_ij_* is a spatial mask and *v_k_* a set of feature pooling weights. We fitted models with increasing number of convolutional layers (one to five). We found that optimizing the final nonlinearity, *f* (*x*), of each neuron was important for optimal performance of the data-driven CNN. To do so, we took the following approach: we split *f* (*x*) into two components:

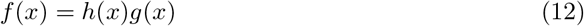

where *g*(*x*) is ELU shifted to the right and up by one unit (to make it non-negative – firing rates are non-negative):

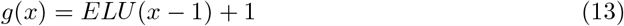

and *h* is a non-negative, piecewise linear function:

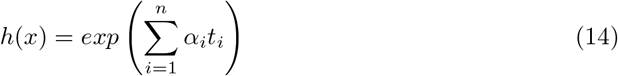

Here, *α_i_* are parameters learned jointly with the remaining weights of the network and the *t_i_* are a set of ‘tent’ basis functions to create a piecewise linear function with interpolation points *x_i_* = −3, − 2.82, …, 6 (i.e. ∆*x* = 0.18):

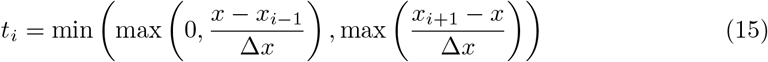

We regularize the output nonlinearity by penalizing the *L*_2_ norm of the first and second discrete finite differences of *α_i_* to encourage *h* to be close to 1 and smooth:

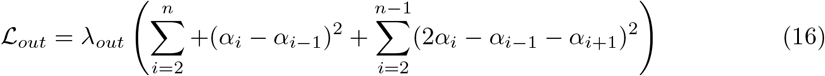

Note that we applied this optimization of the output nonlinearity only to the data-driven model, as doing the same for the VGG-based model it did not improve performance. One potential reason for this difference is that the VGG-based model has a much larger number of feature maps (256 for layer conv3 1) that each neuron can pool from.

### Linear Nonlinear Poisson Model (LNP)

We implemented a simple regularized LNP Model [46]. This model is fitted for each neuron separately and consists of two simple stages: The first one is a linear filter **w** with the same dimensions as the input images. The second is a pointwise exponential function as nonlinearity that converts the filter output into a non-negative spike rate. The LNP assumes spike count generation through a Poisson process, so we minimize a Poisson loss (negative log-likelihood) to obtain the kernels of each neuron (see first term of Equation 17 below). Additionally, we imposed two regularization constraints that we cross-validated: smoothness (Eq 7) and sparsity (Eq. 6). With the same *M* image-response pairs (**x***, y*) of the training set that we used for all other models, we optimized the following loss function:

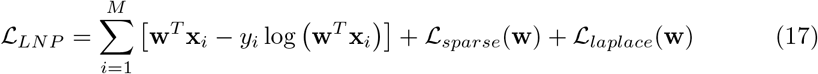

### Gabor Filter Bank Model (GFB)

We implemented a convolutional Gabor filter bank (GFB) model based on the implementation from STRFlab.^1^ Varying versions of the Gabor filter bank model have been used in classical work on system identification [22, 47, 48, 64]. This model convolves quadrature pairs of Gabor filters with varying size, frequency and orientation with the input images that results in an ‘even’ and ‘odd’ linear feature spaces. We considered both a GFB energy model built with the spectral power of each pair (i.e. sum of the squares), and a full GFB model consisting on the GFB energy model appended to the linear even and odd feature spaces.

More specifically, the Gabor filters obeyed the following equations where *x* and *y* represent here both spatial dimensions:

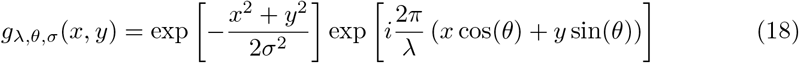

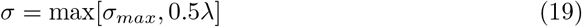

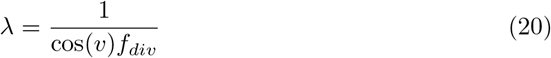

The wavelength (*λ*) of the Gabor function (*g*) is a function of the velocity (*v*) and the frequency *f_div_* of the division (scale). We built the Gabor filter bank using velocities *v_n_* = *nπ*/(2*N_v_*); *n* = 0, …, *N_v_*, orientations *θ_n_* = 2*nπ*/*N_θ_*; *n* = 0, …, *N_θ_*, and frequencies *f_n_* = *n*(*f_max_* − *f_min_*)/*N_scales_* + *f_min_*; *n* = 0, …, *N_scales_*. We chose after cross-validation *N_θ_* = 8, *N_v_* = 2, *N_scales_* = 15, *f_max_* = 9, *f_min_* = 2, and *σ_max_* = 0.3. The kernel size of each Gabor filter was then *k_s_* = *Mσ*/*σ_max_* where *M* is the size of the input image.

The Gabor filter operation is the convolution (⊗) of the image and the Gabor filter. We additionally downsample the resulting feature map of each convolution to *W* × *W*, where *W* is the closest odd integer to *M*/*k_s_* effectively reducing dimensionality. The complex output of this operation (*G_λ,θ_*(*x, y*)) can be decomposed into real (*even*, *E_λ,θ_*(*x, y*)) and imaginary (*odd*, *O_λ,θ_*(*x, y*)) parts. Based on these, we can compute the squared magnitude response *A_λ,θ_*(*x, y*) which is the energy model feature space:

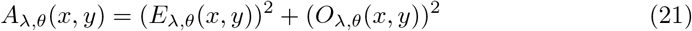

The full feature space of this model consists of 720 feature channels, subdivided into three sets (even, odd, energy) of 240 features each (8 orientations, 2 velocities and 15 scales). On top of this feature space, we then fit an *L*_1_-regularized dense linear readout with an ELU (see Eq. 10) output nonlinearity and a Poisson loss.

### Number of parameters to be learned

The parameters we fit for each of the models belong either to a shared set for all neurons (the core), or are specific to each neuron (the readout). Table 2 shows the number of parameters for each of the models and how many belong to either core or readout. For both the LNP and GFB models, we learn only a readout from a fixed feature space (LNP: one channel of pixel intensities; GFB: 720 channels of wavelet features) for each neuron plus a bias. For the three-layer CNN, we have 32 channels in all layers (32×3 biases) and filters with sizes 13×13×32, 3×3×32×32, and 3×3×32×32, resulting in 23, 963 core parameters. The output feature space for an image is 28××28×32 (reduced from 40×40 due to the padding of the convolutions: no padding in first layer, zero padding in second and third). With a factorized readout and a bias, the readout per neuron is then 28×28 + 32 plus a bias. In addition, our point-wise output nonlinearity has 50 parameters. Thus, overall we have 867 readout parameters per neuron for this CNN model.

For the VGG-based model, although we do not learn the feature space, we do learn batch normalization parameters at the output of the last convolutional layer. For the model that used conv3 1 (256 feature channels) this means learning scale and bias parameters common to all neurons: 2 * 256 = 512 for the core. For a 40×40 input, the output of the feature space is 10×10×256 (due to downsampling twice via max pooling). Here, we learn a dense readout and a bias, so the readout per neuron has 10 × 10 × 256 + 1 = 25, 601 parameters.

### Performance evaluation

We measured the performance of all models with the fraction of explainable variance explained (*FEV*). That is, the ratio between the variance accounted for by the model (variance explained) and the explainable variance (numerator in Eq. 3). The explainable variance is lower than the total variance, because observation noise prevents even a perfect model from accounting for all variance. We compute *FEV* as

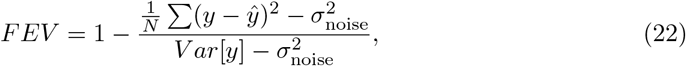

where *ŷ* represents the model predictions, *y* the observed spike counts, and the level of observation noise, 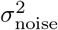 is defined in Eq. 4 above.

### Implementation details

We implemented all models in TensorFlow [65]. We optimized them with Adam [61] using mini-batches of size 256, and early stopping: we evaluated performance on the validation set every 100 training steps, and after ten iterations of no improvement, we decayed the learning rate by a factor of three and repeated this three times. The learning rate at the beginning of the optimization was cross-validated for the goal-driven models and set to 1e-4 for the others as this value always worked best.

#### Tools

We managed our data and kept track of models, parameters, and performance using DataJoint [66]. In addition, we used Numpy/Scipy [67], Matplotlib [68], Seaborn [69], Jupyter [70], Tensorflow [65], and Docker [71].

## Acknowledgments

We thank Philipp Berens and James Cotton for valuable discussions and help during the early stages of this project. We thank Tori Shinn for help with animal training and neural recordings. Research reported in this publication was supported by the German Research Foundation (DFG) grant EC 479/1-1 to A.S.E and the Collaborative Research Center (SFB 1233, Robust Vision); the Bernstein Center for Computational Neuroscience (FKZ 01GQ1002); the German Excellency Initiative through the Centre for Integrative Neuroscience Tübingen (EXC307); the National Eye Institute of the National Institutes of Health under Award Numbers R01EY026927 (A.S.T.), DP1 EY023176 (A.S.T.), and NIH-Pioneer award DP1-OD008301 (A.S.T). The content is solely the responsibility of the authors and does not necessarily represent the official views of the National Institutes of Health. We thank the International Max Planck Research School for Intelligent Systems (IMPRS-IS) for supporting S.A.C. This research was also supported by NEI/NIH Core Grant for Vision Research (EY-002520-37), NEI training grant T32EY00700140 (G.H.D) and F30EY025510 (E.Y.W.). L.A.G was supported by German National Academic Foundation. This research was also supported by Intelligence Advanced Research Projects Activity (IARPA) via Department of Interior/Interior Business Center (DoI/IBC) contract number D16PC00003. The U.S. Government is authorized to reproduce and distribute reprints for Governmental purposes notwithstanding any copyright annotation thereon. Disclaimer: The views and conclusions contained herein are those of the authors and should not be interpreted as necessarily representing the official policies or endorsements, either expressed or implied, of IARPA, DoI/IBC, or the U.S. Government.

## Footnotes

1 strflab.berkeley.edu/

